# Comparative efficiency of eDNA, camera traps and scat surveys to detect a semi-aquatic mammal across multiple catchments

**DOI:** 10.64898/2026.04.28.721338

**Authors:** Simon Lacombe, Sébastien Devillard, Louise D’Hollande, Yann Raulet, Vincent Sablain, Louis Barbu, Geoffrey Didier, Raphaël Mathevet, Claude Miaud, Clément Oyon, Eve le Pommelet, Sébastien Richarte, Serge Rouvière, Alice Valentini, Nathalie Vazzoler-Antoine, Olivier Gimenez

**Affiliations:** CEFE, Université de Montpellier, CNRS, EPHE, IRD, 1919 Route de Mende, 34090 Montpellier, France; Université Claude Bernard Lyon 1, LBBE, UMR 5558, CNRS, VAS, Villeurbanne, 69622, France; EPTB Symbo, boulevard de la Démocratie, F-34130 Mauguio, France; Ville de Montpellier, service Nature Observatoire et Territoire, Direction Paysage et Biodiversité, 2733 avenue Albert Einstein 34000 Montpellier; EPTB Lez (Syndicat du Bassin du Lez – SYBLE), Domaine de Restinclières, F-34730 Prades-le-Lez, France; Association Fiber Nature, 389 rue de l’Espinouse, F-34090 Montpellier, France; EPTB Vidourle, 216 chemin de campagne - 30250 Sommières, France; Métropole de Montpellier, service GEMAPI, 50 place Zeus, CS 39556, 34961 Montpellier, France; SPYGEN, 17 rue du Lac Saint André, CS 20274, F-73375 Le Bourget-du-Lac cedex, France

**Keywords:** Camera-trapping, Environmental DNA, Eurasian Otter, Non-invasive monitoring, Sign-surveys, Wildlife monitoring

## Abstract

Semi-aquatic mammals lie at the intersection of several key conservation issues such as wetland deterioration or species invasions, and monitoring their distribution in space and time is essential to inform conservation strategies. However, gathering information about their presence is challenging due to their elusive lifestyle and generally low abundance. The Eurasian otter (*Lutra lutra*), a near-threatened and strictly protected species in Europe, is currently recolonizing part of its historical range. Its high conservation interest, combined with a dynamic more commonly associated with range-expanding or invasive species, makes it a particularly compelling case study. Otter monitoring has traditionally relied on scat surveys, but recent environmental DNA (eDNA) and camera-trapping initiatives have emerged offering promising complementary tools. Yet, these approaches have rarely been formally compared, either to one another or across regions. Here, we compared the efficiency of spraint surveys, camera traps, and eDNA for detecting otters, and assessed how their performance varied among four catchments in southern France where the species is known to be present. All three methods provided otter detections with varying efficiency. Scat surveys were the most effective method, with an average detection probability of 0.71 and no strong variability between catchments. Although camera-traps had the lowest detection rate, they provided detections at two of the four sites where no spraint was found, highlighting the complementarity of these two approaches. Detection rates varied greatly between individual cameras rather than between catchments, underscoring sensitivity to camera-placement. eDNA showed important variability between catchments, with detection probabilities differing by roughly sixfold across regions. All in all, our results highlight differences in efficiency between methods and across environmental conditions, and show the value of combining approaches for future monitoring programs.

## 1 Introduction

Biodiversity monitoring provides the empirical foundation needed to assess the status of species and ecosystems and to guide management decisions (Nichols and Williams, 2006; Yoccoz et al., 2001). Reliable information on the spatial and temporal dynamics of populations helps identify conservation priorities and objectives (Altermatt et al., 2025; Linden-mayer and Likens, 2009). Monitoring ecological systems also generates the data required to develop and calibrate models that can disentangle complex threats and predict how populations and ecosystems will respond to increasing threats or to conservation interventions (Lindenmayer and Likens, 2009; Nichols and Williams, 2006).

Freshwater ecosystems support a large proportion of continental biodiversity but are particularly vulnerable to global changes (Strayer and Dudgeon, 2010). Among the affected fauna, semi-aquatic mammals is a diverse group lying at the intersection of key conservation issues. Anthropogenic disturbances have led to distributional changes in various semi-aquatic species over the last century (Põdra and Gómez, 2018; Quaglietta et al., 2018; Schertler et al., 2020). For several species, these changes resulted in range contractions and population declines, and many are now considered threatened and targeted by important conservation measures (Põdra and Gómez, 2018; Charbonnel et al., 2016; Quaglietta et al., 2018). By contrast, there are growing examples of invasive semi-aquatic mammals whose ranges are expanding rapidly, threatening native species (Charbonnel et al., 2016; Põdra and Gómez, 2018; Schertler et al., 2020). Furthermore, some native semi-aquatic mammals are acknowledged for their positive effects on ecosystems (Law et al., 2017), as well as their appeal to the general public (Gandy and Watts, 2021; Stevens et al., 2011), making them effective flagship species for stimulating conservation efforts. Collecting data on semi-aquatic mammals presents significant challenges. Many of these species spend much of their time in the water, occur at low densities and are nocturnal, making observations difficult (Hood, 2020). Traditionally, monitoring has relied on indirect indicators (e.g. tracks or scats) of their presence (Bonesi and Macdonald, 2004; Croose et al., 2019; Harrington et al., 2010). This method is generally effective (Bonesi and Macdonald, 2004; Charbonnel et al., 2014) and has been used for mapping the distribution of semi-aquatic species at different scales (Quaglietta et al., 2018; Lacombe et al., 2025; Schertler et al., 2020). However, sign surveys are time-consuming, require substantial field effort and physical access to riverbanks which is often difficult, particularly along large or densely vegetated waterways. Furthermore, accurate identification of signs requires taxonomic expertise or molecular tools (Harrington et al., 2010; Quaglietta et al., 2018). Finally, for many species, availability of scats depends on marking behaviors, which can be affected by external factors like food availability or intraspecific competition (Sittenthaler et al., 2020).

In recent years, camera traps (CTs) and environmental DNA (eDNA) from river samples have emerged as alternative monitoring methods that can help address these limitations. Both are cost-effective and relatively practical to implement in the field (Broadhurst et al., 2021; Burgher et al., 2024; Croose et al., 2023). CTs can be deployed for extended periods with minimal maintenance while eDNA is relatively easy to collect (Broadhurst et al., 2021; Buxton et al., 2022) and can be scaled up across large areas even from a limited number of sites thanks to the fractal nature of rivers and the downstream transport of DNA (Deiner et al., 2016; Lyet et al., 2021). These approaches have proven effective in detecting species that are otherwise difficult to monitor, particularly those that are elusive, occur at low densities, or leave few detectable signs (Harper et al., 2019b; Holm et al., 2023). Notably, both CTs and eDNA have contributed to the early detection of invasive species (Sales et al., 2020; Takaba et al., 2024) and the rediscovery of species considered locally extinct (Croose et al., 2023; Holm et al., 2023) or in the process of natural recolonization (Ballini et al., 2024). However, their effectiveness for monitoring semi-aquatic mammals remains variable. Rare and elusive species tend to have low encounter rates on CTs (Findlay et al., 2023; Lerone et al., 2015), often resulting in low detection probabilities (Sales et al., 2020; Sanders et al., 2024). Moreover, detection rates are sensitive to trap placement, which can introduce variability unrelated to species presence or abundance (Burton et al., 2015; Harper et al., 2019b). Similarly, early applications of eDNA to semi-aquatic mammals reported limited success, with overall low detection probabilities (Harper et al., 2019b; Iacaruso et al., 2025; Sales et al., 2020; Thomsen et al., 2012). Detection rates appear to depend on species ecology, with carnivorous species showing much lower detectability than herbivorous species occurring at higher densities and with a stronger dependency on water (Burgher et al., 2024; Harper et al., 2019b; Sales et al., 2020). Still, application of eDNA to semi-aquatic biodiversity is rapidly evolving and recent studies suggest increasing ability to detect semiaquatic animals including carnivores (Croose et al., 2023; Girolamo et al., 2024; Jamwal et al., 2023). Although many studies provided formal comparisons of these approaches their findings often diverged, suggesting that the relative performance of monitoring methods may be dependent on environmental factors. Despite that, little attention has been given to how variable these methods perform across different ecological contexts.

In this study, we aim to compare the efficacy of sign surveys, camera traps and eDNA for detecting the Eurasian otter (*Lutra lutra*) across four catchments in Southern France – the Lez, Mosson and Vidourle Rivers and the Étang de l’Or pond and river system. We also assess the extent of variability in detection rates both within and between catchments, to evaluate the consistency and robustness of each method under differing ecological conditions. The otter is a particularly compelling study species: it is a near threatened species strictly protected in Europe (Loy et al., 2022), showing a strong conservation interest. At the same time, it is currently recolonizing in many European countries raising monitoring questions typically associated with range-expanding carnivores or invasive species. An otter monitoring protocol based on spraints (otter scats) (Reuther et al., 2000) is being used across several European countries (Kuhn et al., 2019; Loy et al., 2010). In addition, some eDNA and CTs initiatives have appeared efficient to detect the species (Holm et al., 2023; Jamwal et al., 2023; Sales et al., 2020), but always with few sites over relatively small areas, and these approaches have, to our knowledge, never been formally compared. Here, we predict that sign surveys will consistently outperform other approaches, as marking behavior plays a central role in otter ecology (Almeida et al., 2012; Sittenthaler et al., 2020). We also expect detectability with eDNA and CTs to exhibit significant between-catchment variability, with higher detection rates in catchments that have been recolonized by the species for a longer period. Finally, we predict that sensitivity to camera placement will cause stronger within-catchment variability in detectability with CTs compared with other methods.

## 2 Material and methods

### 2.1 Study area

The study took place in southern-France, in the Gard and Hérault departments. We focused on four river systems: the Lez River, the Mosson River, the Étang de l’Or pond and river system, and the Vidourle River (Fig. 1). These systems share a Mediterranean climate, characterized by hot, dry summers and intense rainfall in autumn, which can cause rapid changes in river discharge and intermittent flow in smaller tributaries. The riparian vegetation across these catchments is similar, reflecting typical Mediterranean lowland river communities. These rivers are all subject to engineering to varying degrees, with numerous artificial structures altering their natural flow and morphology along much of their course.

**Figure 1:**
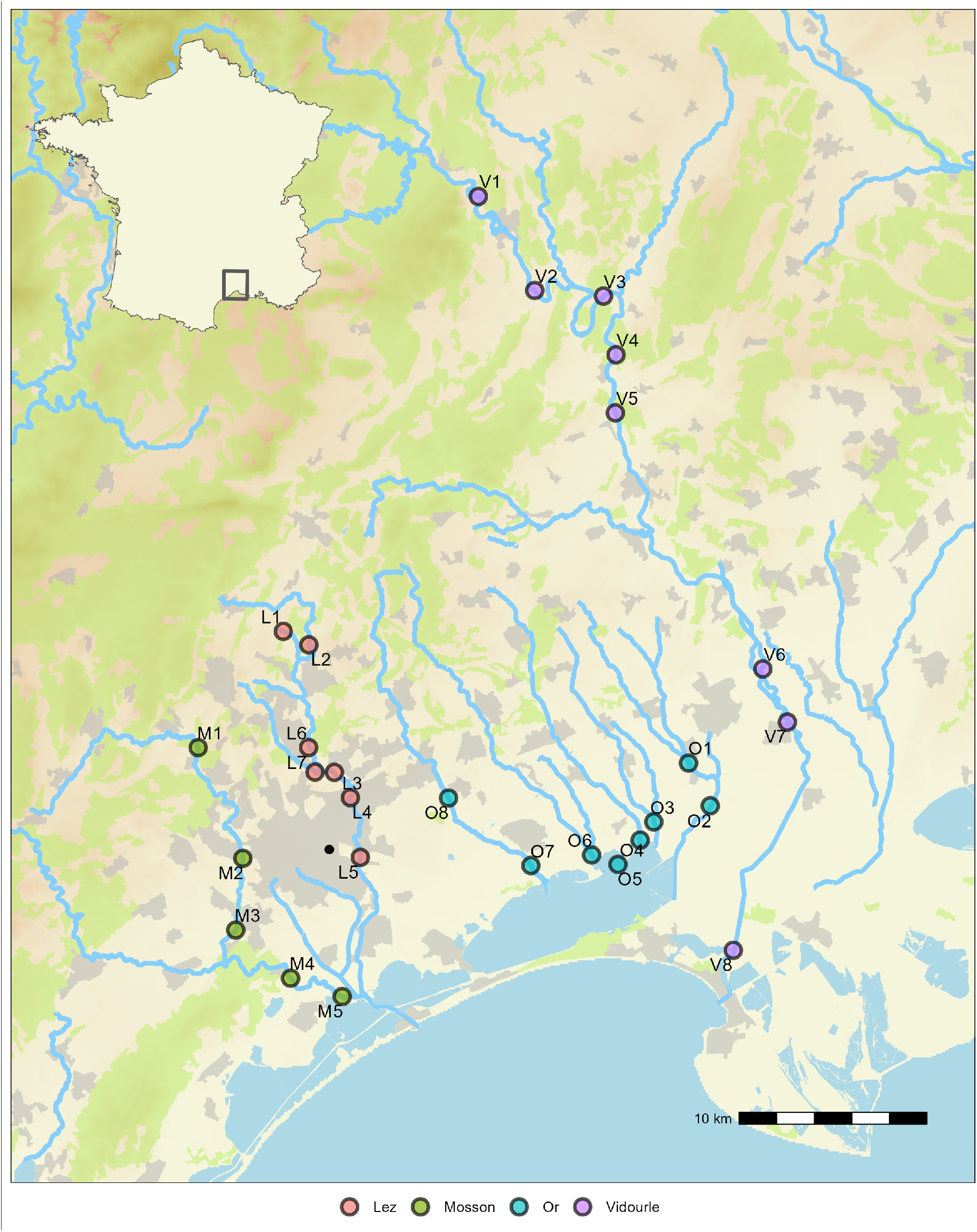
Map of the study area. Sites are shown with color circles, with color indicating the catchment. Rivers, coastal ponds and the sea are shown in blue, forested areas are shown in green, urban areas are shown in grey, and altitude is shown with shades of beige, with darker tones indicating higher altitudes. The black dot shows the city of Montpellier.

The Lez and Mosson rivers both cross the Montpellier metropolitan area, flowing through heavily urbanized landscapes, with intermittent sections of more natural corridors with variable riparian conditions. The Lez is the most heavily engineered river of the study area, featuring numerous artificial weirs and bridges along its 29.6 km course, especially in the upstream part. The Étang de l’Or system comprises a coastal lagoon connected to the Mediterranean Sea, several rivers and artificial canals, as well as extensive marshes and wetlands. The watershed encompasses a variety of landscapes, ranging from scrubland in the upstream section to a mosaic of agricultural, urban, and wetland habitats towards the lagoon. The Vidourle River flows from the Cévennes foothills to the Mediterranean Sea. In its upper half, it flows through relatively natural lands featuring forested areas with dense riparian woodlands and a low level of human disturbances. The rest of the river’s course is more typical of a lowland coastal river with largely rural and agricultural catchment. In this section, the river also passes through several small to medium-sized towns. All four systems have been recolonized by the otter in the last two decades (Lacombe et al., 2025). First sightings date back to 2016, 2008, 2019 and 2012 for the Lez, Mosson, Étang de l’Or and Vidourle respectively. Since then, opportunistic sightings are reported along the course of these rivers with increasing frequencies. To our knowledge, the Vidourle is the only river in the study area in which otters are regularly observed during the day.

### 2.2 Study design

For each catchment, fieldwork was conducted over a two-month period, either in late autumn (Lez in 2023 and Vidourle in 2024) or early spring (Or in 2024 and Mosson in 2025). Therefore, data collection overlapped temporarily for all sampling methods within a catchment, but did not overlap between catchments. We selected 5 to 8 sites per catchment, defined as 600m long river stretches (Fig. 2A). This resulted in a total of 28 sites. Sites were chosen to ensure convenient access to the river and to allow as much of the 600 m stretch as possible to be covered by wading. A minimum distance of 1 km was set between sites to reduce spatial correlation.

**Figure 2:**
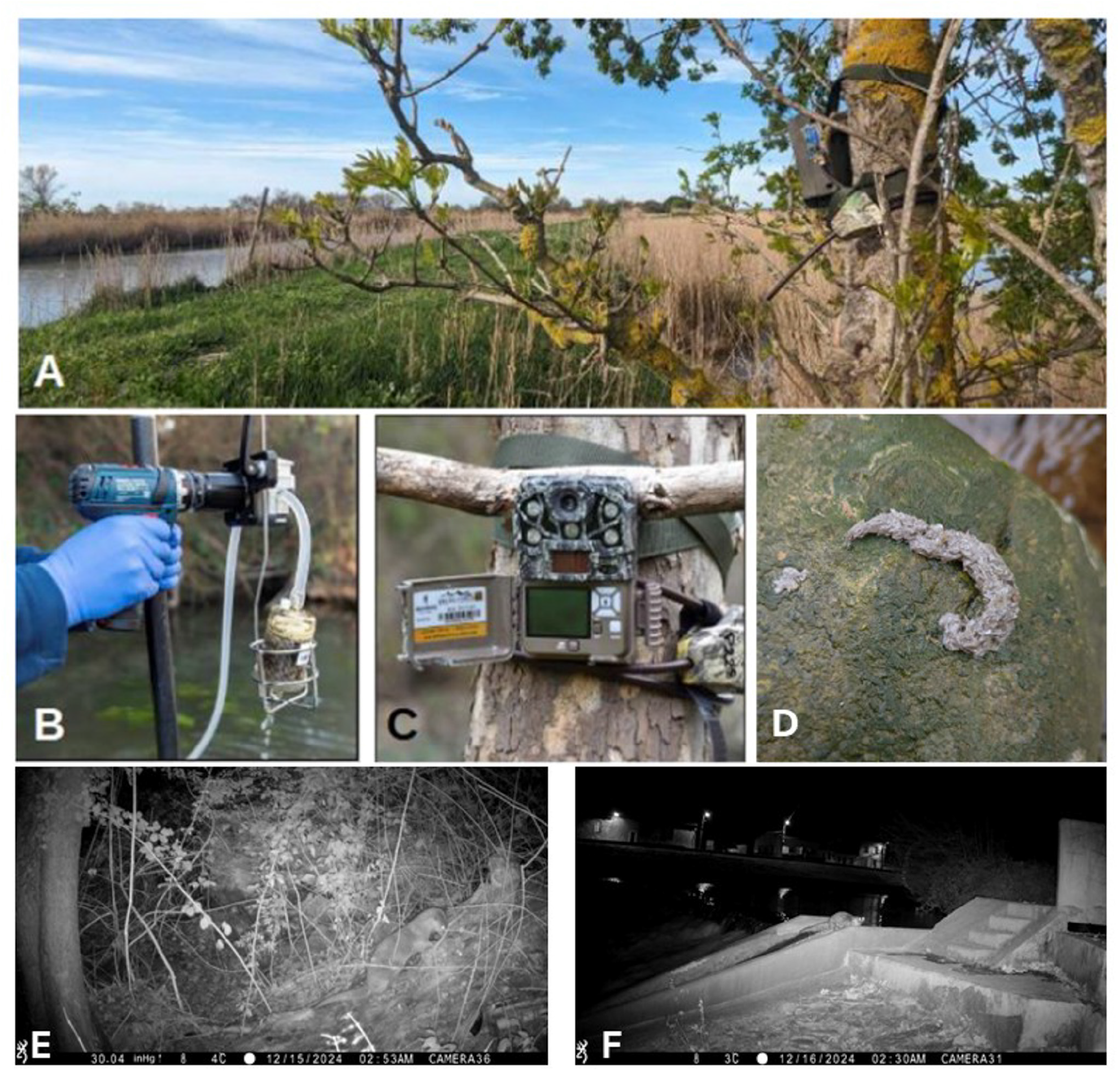
Overview of the monitoring design used to detect the Eurasian otter. A. View of site O2 showing the downstream camera trap and a portion of the river; B. Picture of the peristaltic pump and filtration capsule used to take eDNA samples; C. One of the camera traps used in the study D. An otter spraint. E. F., Eurasian otter marking (E.) and crossing a weir (F.) recorded by two camera traps.

### 2.3 Sign surveys

Each site was surveyed three times during the study period. At each visit, we prospected the river using a protocol adapted from the standard protocol for otter monitoring (Kuhn et al., 2019; Reuther et al., 2000). The latter consists of covering 600m of river along one bank, looking for scats. We slightly modified this protocol by covering the two banks whenever the river could be crossed by foot, and by thoroughly surveying all engineering structures (e.g., weirs, bridges) present on the site. This was done to compensate for the fact that a significant proportion of our sites could not be covered over the entire 600m. Although otter spraints are relatively easy to identify due to their appearance (Fig. 2D), content and very characteristic smell, a risk of misidentification remained (Harrington et al., 2010). We therefore made sure that at least one of four observers trained at recognizing otter spraints was present at each prospection.

### 2.4 Camera trapping

We set up two camera traps (Browning SPEC OPS ELITE HP5, Fig. 2A & 2C) at each site: one close to the upstream part of the site and one close to the downstream part. Where possible, we positioned the cameras to focus on microhabitats frequently used by otters, such as small paths leading from the river into the riparian zone, wooden stumps, artificial structures like bridges or weirs, or gravelly banks or islets that are often used to consume certain preys (Kim and Hong, 2024; Pagh et al., 2025). Cameras were set to operate in motion-sensor recording 30s long videos at trigger. Cameras were active for around two months, and revisited regularly to change batteries and SD cards. Since some cameras remained in place beyond the end of the study period, we only considered the first 77 days of observation. This corresponds to the deployment duration in the Vidourle catchment, which had the shortest camera deployment time. All videos were viewed and species identified by the first author (**SL**). We then aggregated the data into 24-hour periods starting at midday, providing daily detection/non-detection histories for all camera traps.

### 2.5 eDNA sampling

Each site was also visited once to collect water samples for eDNA analyses. Samples were collected near the downstream end of the site, in the main course of the river. For each sample, we filtered the river water for 30 minutes (accounting for c.a. 30L) using a peristaltic pump (Vampire sampler, Burkle, Germany; Fig. 2B) and disposable sterile tubes into VigiDNA 0.45µM crossflow filtration capsules (SPYGEN, le Bourget du Lac, France; Fig. 2B). At the end of the filtration, the cartridge was drained of water and filled with 80 mL of CL1 preservation buffer (SPYGEN). This procedure was repeated three times at the same location, accounting for a total of three eDNA samples per site. The three samples were collected on the same day to ensure similar conditions for a given site. eDNA samples were stored in a box at room temperature until the DNA extraction.

### 2.6 eDNA lab and bioinformatics

The eDNA analysis was performed following methods described in (Pont et al., 2018) using a vertebrate primer (V05, Riaz et al. (2011)) for samples from the Vidourle and Mosson rivers and a mammal primer (Mamm01, Taberlet et al. (2018)) for samples from the Lez and Bassin de l’Or, both located on the 12S region. We used these primers instead of a primer specific to the European otter to align with the monitoring objectives of our local partners – municipalities and watershed agencies – who aimed to survey multiple taxonomic groups. After the DNA extraction, the samples were tested for inhibition by quantitative polymerase chain reaction (qPCR) following the protocol in (Biggs et al., 2015). If the sample was considered inhibited it was diluted fivefold before the amplification. DNA amplifications were performed in a final volume of 25 *µL*, using 3 *µL* of DNA extract as the template and either vertebrate (V05) or mammal (Mamm01) primers. The amplification mixture contained 1 U of AmpliTaq Gold DNA Polymerase (Applied Biosystems, Foster City, California, USA), 10 *mmol*.*L*^*−*1^ Tris-HCl, 5*mmol*.*L*^*−*1^ KCl, 2.5 *mmol*.*L*^*−*1^ MgCl2, 0.2 *mmol*.*L*^*−*1^ each dNTP, 0.2 *µmol*.*L*^*−*1^ of each primer pair, 4 *µmol*.*L*^*−*1^ human blocking primer (De Barba et al., 2014; Lyet et al., 2021) and 0.2 *µg*.*µL*^*−*1^ bovine serum albumin (BSA, Roche Diagnostic, Basel, Switzer-land). The primers were 5’-labeled with an eight-nucleotide tag unique to each sample (with at least three differences between any pair of tags), allowing the assignment of each sequence to the corresponding sample during sequence analysis. The PCR mixture was denatured at 95°C for 10 min, followed by 50 cycles of 30s at 95°C, 30 s at 55 °C (V05) or 57°C (Mamm01), 1 min at 72°C, and a final elongation step at 72°C for 7 min in a room that is dedicated to amplified DNA and has negative air pressure and physical separation from the DNA extraction rooms (with positive air pressure). Twelve replicate PCRs were run per filtration capsule. The purified PCR products were pooled in equal volumes to achieve a theoretical sequencing depth of 300,000 reads per sample.

The sequencing libraries were prepared using the TruSeq kit (Illumina) following the manufacturer’s instructions and then they were sequenced on a NextSeq sequencer using P1 XLEAP-SBS Reagents. Extraction and amplification negative controls were performed for each batch of extraction and amplification, and they were sequenced in parallel of the samples (12 amplification per negative control). Subsequently, bioinformatic analysis and taxonomic assignment of sequences were conducted following the methodology outlined in (Valentini et al., 2016). In summary, the sequence reads underwent analysis utilizing the OBITools package (Boyer et al., 2016). Initially, the forward and reverse reads were merged and then demultiplexed and dereplicated. Each sample was then segregated into a distinct dataset by partitioning the original dataset into multiple files. Sequences shorter than 20 bp, occurring less than 10 times per sample, or identified as “internal” by the obiclean program were excluded. The program ecotag was used for the taxonomic assignment of sequence with the sequence retrieved from the public reference database GenBank (version 247). Only sequences with a similarity of higher than 98 % were kept. Considering the bad assignments of a few sequences to the wrong sample due to tag-jumps (Schnell et al., 2015), all sequences with a frequency of occurrence below 0.001 per taxon and per library were discarded. Subsequently, the obtained data underwent curation for Index-Hopping (MacConaill et al., 2018). For this purpose, a threshold was empirically determined for each sequencing batch using experimental blanks, which consisted of tag combinations not present in the libraries for a specific sequencing batch. Species found in the negative controls included only human and domestic animals.

### 2.7 Statistical analysis

We estimated otter detection probabilities by fitting generalized linear mixed models (GLMM) to the detection/non-detection data, with binomial distributions and logit link functions. We fitted three separate GLMMs, one for each monitoring method, for estimating an average detection probability without fixed effect and with a catchment random intercept to quantify regional variability. For the camera-trap data, we also used a camera random effect to capture variability due to camera placement. For camera trapping, we estimated daily detection probabilities since cameras’ activity periods were highly variable. To facilitate comparison between methods, we reported the CT detection probability for 30 days, calculated as 1 *−* (1 *− P*_*CT*_)^30^, where *P*_*CT*_ is the estimated detection probability for one day. We also calculated detection curves for each method, giving the probability to detect the species as a function of the number of replicates.

We fitted the models within the Bayesian framework using R version 4.4.2 (RCoreTeam, 2026) and the R package brms (Bürkner, 2017). Default weakly informative Student-t(3, 0, 2.5) priors were applied to the intercept and random-effect standard deviations, and models were run for 5,000 iterations (1,000 warm-up) across two chains. We assessed convergence by visually inspecting trace plots and ensuring 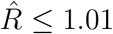 for all parameters.

## 3 Results

### 3.1 Sign surveys

Of the 28 sites in the study area, only four did not have any positive survey for otter signs (L7, O4, O5 and V8, Fig. 3). For all others, the average number of positive surveys was 2.42 [2.09,2.74] (mean + 95% CI). In total, it resulted in 58 positive surveys out of 84 (Fig. 3). The sign surveys estimated detection probability was the highest of all methods (0.71 [0.42,0.91], Fig. 4). The estimated standard deviation for the catchment random effect was low 0.87 [0.06,3.2] (Fig. 5) leading to no clear differences in the detection probability between the four catchments (Fig. 4).

**Figure 3:**
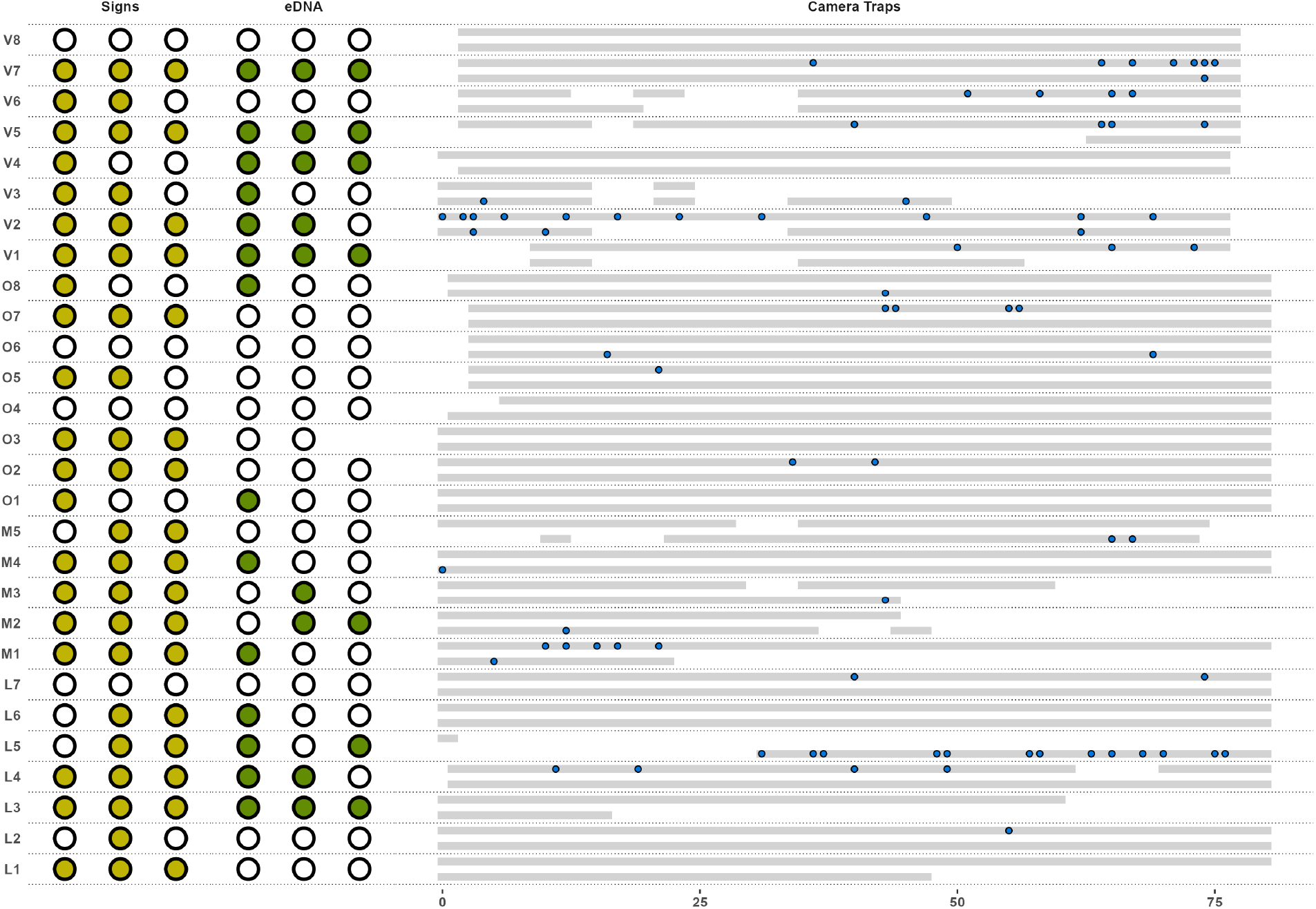
Raw detection data for each site. Empty circles show non-detection for sign sampling (yellow) and eDNA (green). Grey lines show the activation period of each cameratrap, with blue dots showing days with detections of otters.

**Figure 4:**
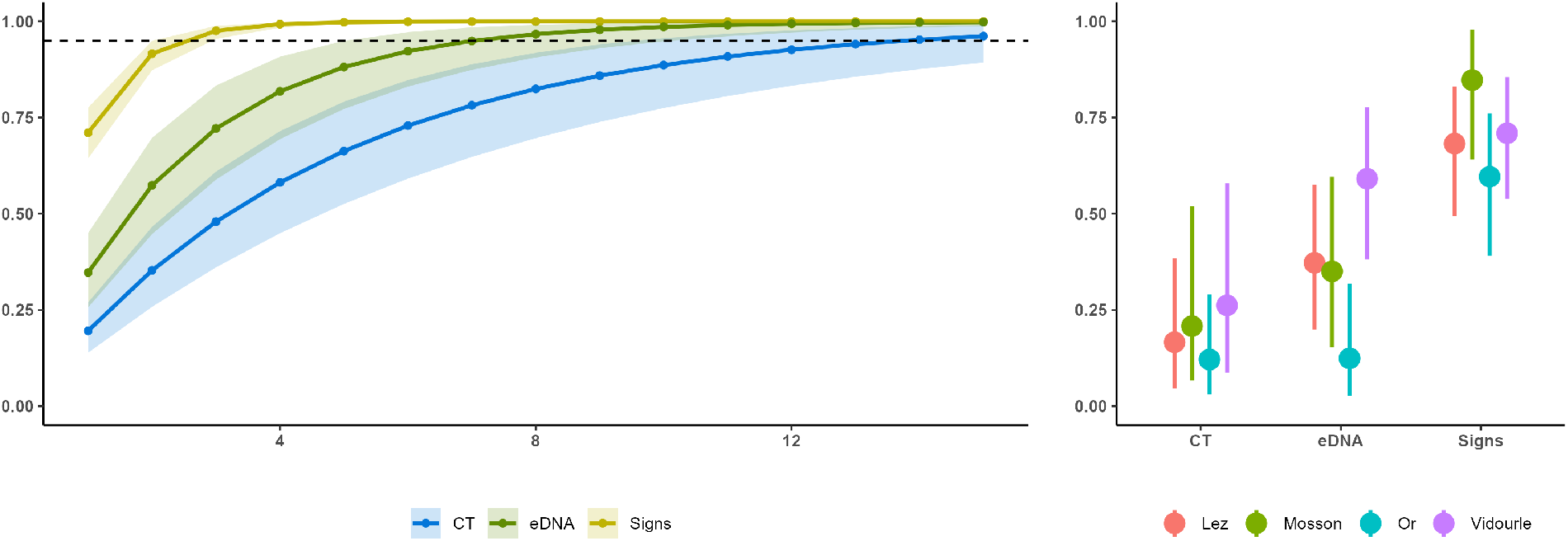
Detection curves (left) and estimated detection probabilities (right). Detection curves show the probability of having at least one detection as a function of the number of field replicates – defined as a camera-trap active for a month, an eDNA sample or a sign-survey. The dashed line shows the 95% probability line. 95% credible intervals are shown with error bars for the detection probabilities. For readability the 50% credible intervals are shown for the detection curves.

**Figure 5:**
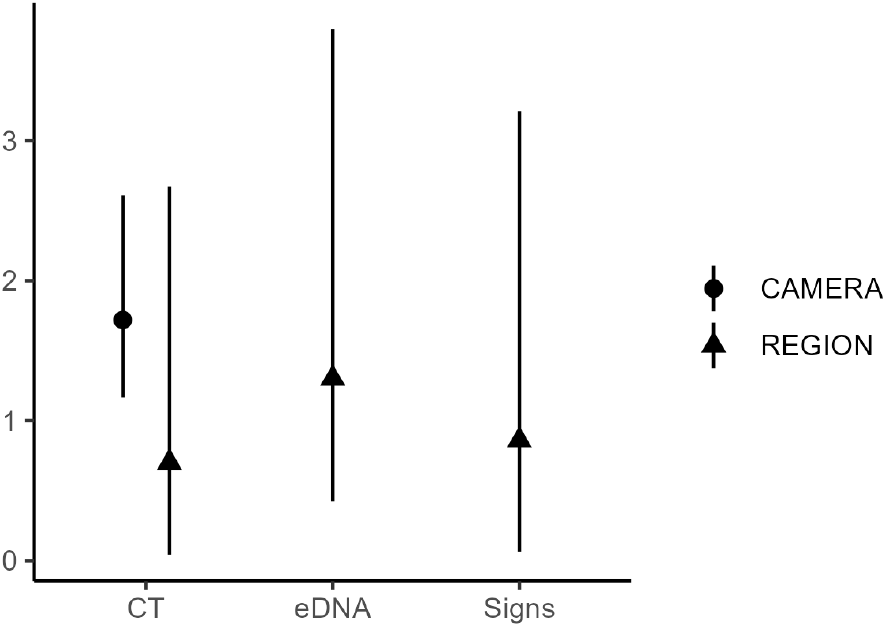
Standard deviation associated with each random effect estimated by the model. Error bars show 95% credible intervals.

### 3.2 Camera trapping

On average, cameras were active for 65.9 [60.3,71.5] days, with a minimum activation period of two days due to camera malfunctioning (L5, Camera 14, Fig. 3). In total, 16 cameras were active for less than 60 days (Lez: 4, Mosson: 6, Vidourle: 6), among which six malfunctioned and were replaced (including four after a major flooding on the Vidourle), eight malfunctioned and were removed (including one after the latter flooding), and two were stolen. During the study period, a total of 26,925 videos were recorded of which 3,153 were attributed to wild mammalian species. Fifteen different species or group of species were identified (Appendix 1, Table. S1). 111 of these videos were of Eurasian otters, accounting for a total of 79 positive camera days at 23 different cameras spanning 20 sites. In particular, we detected otters at two of the four sites where no spraint was found. The estimated detection probability was the lowest of all methods (0.19 [0.06,0.57] for 30 CT-days, Fig. 4). We did not find signs of strong between-catchment variability with a standard deviation for the catchment random effect of 0.68 [0.03,2.64] (Fig. 5) and no clear differences in the estimated detection probability of each catchment (Fig. 4). However, we found a very strong effect of the camera random effect, with a standard deviation of 1.69 [1.14, 2.55] (Fig. 5) and a monthly detection probability ranging from 0.07 [0.002, 0.51] for Camera 25 (V4) to 0.99 [0.94, 1] for Camera 13 (L5) (Fig. S2).

Otter activity was almost exclusively nocturnal, with 93.6 % of all observations occurring between 7 PM and 7 AM. All daily observations occurred in the Vidourle catchment (Fig. S1).

### 3.3 eDNA

Otter DNA was found in water samples for 16 sites out of 28 (57 %) with a total of 30 positive samples out of 83 (Fig. 3). The proportion of positive replicates was highly variable between catchments, going from 8.7% in the Étang de l’Or to 62.5% in the Vidourle (Fig. 3). The average number of detections per site was 1.07 [0.62,1.52], or 1.87 [1.40,2.35] if we only consider sites with at least one detection. The estimated detection probability was 0.34 [0.10,0.73] (Fig. 4). Important variability between catchments is shown by the standard deviation of the catchment random effect (1.28 [0.44,3.52], Fig. 5) and the estimated catchment scale detection probabilities ranging from 0.12 [0.03,0.30] in the Étang de l’Or to 0.59 [0.39,0.78] in the Vidourle (Fig. 4).

## 4 Discussion

In this study, we aimed to evaluate and compare the efficiency of sign-surveys, cameratraps (CTs) and environmental DNA (eDNA) to detect the Eurasian otter in four Mediterranean catchments. Although sites provided spatial replicates for each monitoring method, additional replication differed among approaches: spraint surveys involved temporally separated visits, eDNA was based on multiple field replicates collected on a single day, and camera-trapping relied on two cameras continuously recording detections at each site. These differences may influence detection estimates and should be considered when interpreting comparisons. However, since these differences were accounted for in the analyses and reflect standard replication structures associated with each method, we consider that they do not prevent meaningful comparison.

All methods enabled us to detect the species at a significant proportion of the surveyed sites. Consistently with our first hypothesis, we found that sign-surveys were the most effective method, providing detections at 24 of our 28 sites (Fig. 3). eDNA showed intermediate efficacy with significant variability observed between catchments (Fig. 5). Regarding CTs, important variability in the detection rate was related to a camera effect (Fig. 5), suggesting that fine-scale trap placement is more important than large-scale environmental conditions to detect the species.

### 4.1 Contrasting efficiency of spraint surveys, eDNA and camera-trapping

Otters have high metabolic rates, consume large quantities of food every day (Kruuk, 2006), and have a relatively fast transit, leading them to produce many spraints daily throughout their home ranges (Mirone et al., 2024). Furthermore, otters use scent marking for intra-specific communication (Hutchings and White, 2001; Kruuk, 2006), and they generally deposit spraints in conspicuous and stereotypical places such as rocks, roots or concrete blocks to facilitate smell dispersion (Almeida et al., 2012). Therefore, in places where otter populations are well established, availability of spraints is generally high (Sittenthaler et al., 2020) and they are very easy to locate for trained observers, explaining the high performances of spraint surveys (Fig. 4). Average performances of eDNA were promising, with detections at c.a. 60% of all sites (Fig. 3). However, the important number of sites with just one or two detections suggests that, even when DNA is available in the environment, detectability remains uncertain. We therefore advocate for the systematic use of field replicates to increase the reliability of species presence assessments (Griffiths et al., 2025). Our metabarcoding approach could have lowered the detection efficiency because of competition with sequences from hyperabundant species. However, this effect appeared minimal in our study (see also Condachou et al. (2024)), as high sequencing depths allowed detection of otter DNA even in samples dominated by tens of thousands of coypus reads (Appendix 3). We found that eDNA detection rates were highly variable between regions (Fig. 4, 5). Notably, detection probabilities were around six times higher in the Vidourle than in the Étang de l’Or. eDNA detection rates have been shown to increase with animal densities and activity levels (Baldigo et al., 2017; Thomsen et al., 2012), and many species have lower detection rates in lentic systems due to patchy habitats (Harper et al., 2019a), increased degradation rates and limited DNA dispersion (Burgher et al., 2024). Finally, rainfall prior to sampling is known to enhance detection by flushing DNA into the water (Condachou et al., 2025; Lyet et al., 2021; Macher et al., 2023). In our case, it is difficult to link the observed variability with environmental heterogeneity alone, as regions were sampled in different years and different seasons, with contrasting temperature and rainfall, known to affect eDNA transportation and degradation (Harper et al., 2019a). We also used two different primer sets, which potentially affected detection efficiency. However, primer choice is unlikely to fully explain the observed patterns, as similar detection probabilities were observed in the Lez and Mosson despite the use of different primers. It is more likely that variability instead reflects a combination of hydrology, rainfall and otter densities. Indeed, in the Étang de l’Or catchment, sampling took place during a particularly dry period, and most sites were strongly lentic, potentially contributing to the low detection probability. On the other hand, otters have been present in the Vidourle for longer, and the area is more rural and less disturbed by human activity, likely supporting higher densities and activity and therefore higher eDNA detection rates. Still, our design did not allow formal identification of the drivers of the variability in detection rates, and these explanations should be regarded as hypotheses.

The CT detection probability was lower overall than for the other methods (Fig. 4). Nevertheless, camera traps detected otters at more sites than eDNA, including two sites where no detections were obtained with the other methods. Contrary to eDNA, the performance of CTs did not vary strongly between catchments. Instead, we found that detection varied between individual cameras (Fig. 5, S2) suggesting a high sensitivity to trap placement, as found in previous studies for otters (Gil-Sánchez and Antorán-Pilar, 2020; Harper et al., 2019b) and other species (Burton et al., 2015; Croose et al., 2025; Sanders et al., 2024). This sensitivity is particularly pronounced for semi-aquatic species as their low thermal contrast with the environment reduces the likelihood of triggering infrared sensors (Findlay et al., 2020; Lerone et al., 2015). As a result, otters are less likely to be detected when swimming past a camera, and efficient detection may therefore require targeting locations where they leave the water for e.g. marking, eating prey or overcoming obstacles to river flow (Fig. 2E, F). Previous studies have found that detection rates are improved when cameras are placed near potential latrines (Wagnon and Serfass, 2016), and we suggest that targeting movement bottlenecks – including artificial structures like weirs and dams, or clear paths within the riparian buffer – as well as activity hotspots – including river islets, small gravel beds and shallow areas with emergent vegetated hummocks – is also effective (Appendix 5), as these features constrain movement or increase the likelihood of terrestrial activity.

### 4.2 Complementarity of these monitoring approaches

Although we aimed to sample catchments with contrasting ecological conditions, all surveyed catchments were strongly Mediterranean and therefore represented only a small portion of Eurasian otter habitats. In areas with deeper or less productive rivers (Almeida et al., 2012; Weinberger et al., 2016), or with banks less prone to marking (Loy et al., 2009), spraint surveys could be less effective to assess otter presence. Similarly, the rivers we sampled hosted well-established otter populations, and spraint surveys may be less effective at the colonization front, where populations consist of a few dispersing individuals (Hutchings and White, 2001; Sittenthaler et al., 2020). Actually, practitioners involved in otter monitoring in France have reported difficulties in finding spraints along newly colonized rivers, despite otter presence being asserted by other methods (N. Fuento, pers. comm., 2024). Spraint sampling was also the most time-consuming and impractical method to implement. Even in these Mediterranean rivers, with many shallow sections and accessible banks, it was impossible to sample the entire length of the transect for a majority of sites, and it is likely that in wider, deeper lowland rivers, prospecting would be very difficult if not impossible. Here, we suggest that both eDNA and CTs have the potential to address these two limitations. Both methods required less time in the field (see Appendix 4 for a summary of the costs and labor times involved), and were achievable with only punctual access to the river, showing their potential to increase the range of sites sampled. This is particularly true for eDNA, for which sampling can be done regardless of river morphology or bank-side cover (Altermatt et al., 2025; Beng and Corlett, 2020). Previous studies highlighted CTs and eDNA as capable of providing detections of semi-aquatic mammals where traditional sign surveys fail, in places where individuals are rare or where habitat characteristics hinder the detectability of spraints (Girolamo et al., 2024; Jamwal et al., 2021). We find support for this idea for camera-trapping only. Indeed, CTs yielded detections at two of the four sites where no spraints were found (L7 and O6, Fig. 3), as well as one site where only a single spraint was found over the three visits (L2, Fig. 3). L2 and L7 were located on two tributaries of the Lez with low water levels at the moment of the survey and likely limited food availability (Appendix 5). In such contexts, otters may explore opportunistically without visibly marking the area. At O6, by contrast, cameras yielded two instances of foraging activity (Appendix 5), but dense vegetation and absence of rocks or roots along the riverbanks may have hindered both spraint deposition and detectability. This suggests that otters use certain areas in ways that do not leave detectable signs – either because individuals are only passing through or because the site is unsuitable for marking.

Finally, all three methods can provide different and complementary information. For instance, CTs and eDNA provided information about the community (Table S1 & S2, Appendix 5). CTs also provided information about species activity patterns, behavior or breeding status. In particular, they revealed that otters were strongly nocturnal in the study area (Fig. S1, Gimenez et al. (2025)), confirming a known tendency in human dominated landscapes. Interestingly, a few detections occurred during the day in the upstream part of the Vidourle (Appendix 5), suggesting that diurnal activity may be higher in places with lower human activity, as found in (Penteriani et al., 2025). Spraints can provide information on individual diet (Mirone et al., 2024), be sequenced to infer geographic origin (Pigneur, 2018), or be used to determine individual identity for demographic or socio-spatial analyses.

### 4.3 Conservation and management implications

Coordinated otter monitoring has long relied exclusively on spraint monitoring (Kuhn et al., 2019; Loy et al., 2010). This method prevents sampling habitats that are unsuitable for spraint surveys, and can delay otter detection by several years after establishment due to limited marking at low densities (Sittenthaler et al., 2020). However, early otter detection is particularly important to track the spread of the population, to take timely conservation decisions supporting the establishment of the species in areas where populations are fragile, and to anticipate potential conflicts. Collecting occurrence data where the species struggles to establish is also critical for adaptive monitoring, as it helps identify the drivers limiting populations and supports more informed conservation planning (Lindenmayer and Likens, 2009). Broadening the range of approaches available to practitioners allows sampling a wider range of habitats, and can improve detection efficiency. CTs may be especially valuable in that context, as they can detect individuals even when marking activity is limited, thereby improving the likelihood of early detection. In practice, detection success can be maximized by placing cameras at locations where otters are likely to leave the water – such as riparian wildlife trails, obstacles, or marking sites – highlighting the importance of species-specific ecological knowledge when designing CT surveys. Although we did not find eDNA to increase the number of sites with otter detection, repeating the present study in colonization fronts would allow to further investigate the complementarity of these approaches. We also encourage further research into the environmental and methodological drivers of eDNA detectability for semi-aquatic mammals. Although we focused on Eurasian otters, our findings are applicable to other semi-aquatic mammals, including declining species that urgently need conservation and mostly occur in small densities such as the European mink (Croose et al., 2023) and the Iberian desman (Quaglietta et al., 2018), as well as invasive species like the coypu, for which early detection is critical for management (Coster, 2024; Viviano et al., 2025). Since eDNA and CT analyses can be scaled up to a community level with few additional cost or field effort, this makes them particularly relevant for biodiversity assessments in a context with limited resources. Finally, such studies foster collaborations beyond academic research, creating an effective framework to connect scientists, managers, and stakeholders (Gimenez et al., 2025).

## Acknowledgements

This research benefited from discussions conducted within the DISCAR working group, funded by the French Foundation for Research on Biodiversity (FRB) through its synthesis center CESAB. We thank all the people who supported us in the field, in particular Lucas Buffan, Matthieu Rigal, Axelle Scamps, Tatiana Tronel and the members of the HAIR team at CEFE. We would also like to thank Alessandro Balestrieri and an anonymous reviewer as well as the Editor Vincenzo Penteriani for their review and their constructive comments, which contributed to the improvement of the manuscript. We thank for their financial support the City of Montpellier and Montpellier Méditerranée Métropole through the partnership agreement with the Centre d’Écologie Fonctionnelle et Évolutive (CEFE), the University of Montpellier through its Labex Cemeb, the OSU OREME and the Beauval Nature association. This research was also partly funded by Biodiversa+, the European Biodiversity Partnership, in the context of the Big Picture project under the 2022-2023 BiodivMon joint call. It is co-funded by the European Commission (GA No. 101052342) and the French Agence Nationale de la Recherche (ANR). This research was also partly funded by the ANR through the project nachos for “Interdisciplinary approach to small carnivores - humans relationships” (grant ANR-25-CE03-5469).

## Supplementary Material

Supplementary material, including figures and tables (Appendix 1), codes and data (Appendix 2), results of eDNA samples with the number of reads per detected taxa in each sample (Appendix 3), a summary of the costs and labor times associated with each method (Appendix 4) and a collection of camera-trap videos (Appendix 5) are available at https://sdrive.cnrs.fr/s/cXJ9N9MqyKyTFKy with the password “Petites-loutres2026!”.

### CRediT authorship contribution statement

**Simon Lacombe**: Conceptualization, Data curation, Formal analysis, Investigation, Methodology, Visualization, Writing – original draft, Writing – review and editing. **Sébastien Devillard**: Conceptualization, Methodology, Supervision, Writing – review and editing. **Louise D’Hollande, Alice Valentini**: Data curation, Investigation, Writing — review and editing. **Yann Raulet, Vincent Sablain, Louis Barbu**: Conceptualization, Investigation, Methodology, Writing – review and editing. **Geoffrey Didier, Clément Oyon, Sege Rouviè re, Nathalie Vazzoler-Antoine**: Investigation, Methodology, Writing — review and editing. **Raphaël Mathevet, Claude Miaud**: Conceptualization, Investigation, Validation, Writing - Review and Editing. **Eve le Pommelet, Sébastien Richarte**: Funding acquisition, Investigation, Methodology, Investigation, Writing — review and editing. **Olivier Gimenez**: Conceptualization, Funding acquisition, Investigation, Methodology, Project administration, Supervision, Validation, Writing — review and editing.

### Declaration of generative AI and AI-assisted technologies in the writing process

During the preparation of this work, the authors used ChatGPT to polish the text and enhance the English language. After using this tool, the authors reviewed and edited the content as needed and take full responsibility for the content of the published article.

